# PRDM9 forms a multiprotein complex tethering recombination hotspots to the chromosomal axis

**DOI:** 10.1101/056713

**Authors:** Emil D. Parvanov, Hui Tian, Timothy Billings, Ruth L. Saxl, Catrina Spruce, Rakesh Aithal, Lumir Krejci, Kenneth Paigen, Petko M. Petkov

## Abstract

In mammals, meiotic recombination occurs at 1-2 kb genomic regions termed hotspots, whose positions and activities are determined by PRDM9, a DNA-binding histone methyltransferase. We now show that the KRAB domain of PRDM9 forms complexes with additional proteins to allow hotspots to proceed into the next phase of recombination. By a combination of yeast-two hybrid assay, *in vitro* binding, and co-immunoprecipitation from mouse spermatocytes, we identified four proteins that directly interact with PRDM9’s KRAB domain, namely CXXC1, EWSR1, EHMT2, and CDYL. These proteins are co-expressed in spermatocytes at the early stages of meiotic prophase I, the limited period when PRDM9 is expressed. We also detected association of PRDM9-bound complexes with the meiotic cohesin REC8 and the synaptonemal complex proteins SYCP3 and SYCP1. Our results suggest a model in which PRDM9-bound hotspot DNA is brought to the chromosomal axis by the action of these proteins, ensuring the proper chromatin and spatial environment for subsequent recombination events.

## INTRODUCTION

Genetic recombination assures the proper segregation of homologous chromosomes at the first meiotic division, preventing aneuploidy. It also plays an important evolutionary role by facilitating the creation of new, favorable combinations of alleles and the removal of deleterious mutations by unlinking them from surrounding sequences. In mammals, as in yeast, higher plants, and birds, recombination occurs at specialized sites along chromosomes known as hotspots (6, 34), typically a kilobase or so in length, separated by tens to hundreds of kilobases that lack recombination.

In mammals, a meiosis-specific protein, PRDM9, first identified by our group and others (5, 32, 35), is the primary determinant of recombination hotspot locations (5, 22, 32, 35, 41). PRDM9 combines domains from two large families of proteins - KRAB- zinc finger (ZnF) (30) and PR domain proteins (18) and is the only protein known to contain all three characteristic domains of these families – a KRAB domain implicated in protein-protein interactions, a PR/SET domain with a histone methytransferase activity, and a ZnF domain for DNA recognition and binding. Recombination begins when the C-terminal ZnF domain of PRDM9 recognizes and binds to hotspot specific DNA sequences (5, 7, 32, 35). The PR/SET domain then locally trimethylates histone H3 on lysine 4 (H3K4me3); this results in rearrangement of the local nucleosome pattern, creating a central nucleosome-depleted region (4) where the double strand breaks (DSBs) required for the exchange of DNA sequences between homologous chromatids occur (10). The extent of trimethylation of local nucleosomes delimits the span over which the final genetic crossovers can take place (4). In the absence of PRDM9, DSBs are formed at other available H3K4me3 sites, mainly promoters, but they cannot be repaired properly and germ cells undergo apoptosis (41).

We now show that the PRDM9 KRAB domain plays a crucial role in binding and recruiting additional proteins into multi-protein complexes that bring hotspots into the next phase of recombination; these proteins include CXXC1, a DNA-binding protein with a CXXC domain found in CpG-binding proteins (23), EWSR1, which binds single stranded RNA and DNA (16, 29, 33), EHMT2, a histone methyltransferase catalyzing formation of H3K9me1,2 and H3K56me1 (46, 47), and CDYL, a methyl reader of H3K9me2/3 (15) and H3K27me3 (53), and a putative histone acetylase (26).

A combination of yeast two-hybrid screens and *in vitro* pull-downs of purified proteins showed that the four proteins directly bind to the N-terminal region of PRDM9 where the KRAB domain is located. We further confirmed the *in vivo* interaction between PRDM9 and EWSR1, EHMT2, and CDYL in early prophase spermatocytes by co-immunoprecipitation (co-IP), and cytological colocalization. In spermatocytes, we also detected association of PRDM9-bound complexes with the meiotic cohesin REC8 and the chromosomal axis/synaptonemal complex proteins SYCP3 and SYCP1.

These results suggest that PRDM9-bound hotspot DNA is brought to the chromosomal axis by interaction with other proteins serving as a link between PRDM9 and cohesins/SC proteins, thereby assuring a proper spatial environment for DSB initiation and repair.

## MATERIALS AND METHODS

### Ethics statement

The animal care rules used by The Jackson Laboratory are compatible with the regulations and standards of the U.S. Department of Agriculture and the National Institutes of Health. The protocols used in this study were approved by the Animal Care and Use Committee of The Jackson Laboratory (Summary #04008). Euthanasia for this study was done by cervical dislocation.

### Constructs

pBAD-Prdm9 was described in (7).

pGBKT7-Prdm9 – whole length Prdm9 ORF from pBAD-Prdm9 was amplified with primers

pR638+pR916 and inserted into pGBKT7 using BamHI-PstI restriction sites.

pGBKT7-Prdm9 KRAB–PR-SET – pR638+pR608.

pGBKT7-Prdm9 KRAB – pR790+pR1382.

pGBKT7-Prdm9 ID–PR-SET – pR1658+ pR1661.

pGBKT7-Prdm9 ID/2 – pR1658+pR1383

pGBKT7-Prdm9 ID – pR1658+pR1659

pGAD-T7- Prdm9 construct was created by restriction excision by EcoRI-SalI from pBAD-Prdm9 and insertion by same sites in pGAD-T7 empty vector.

### Antibodies

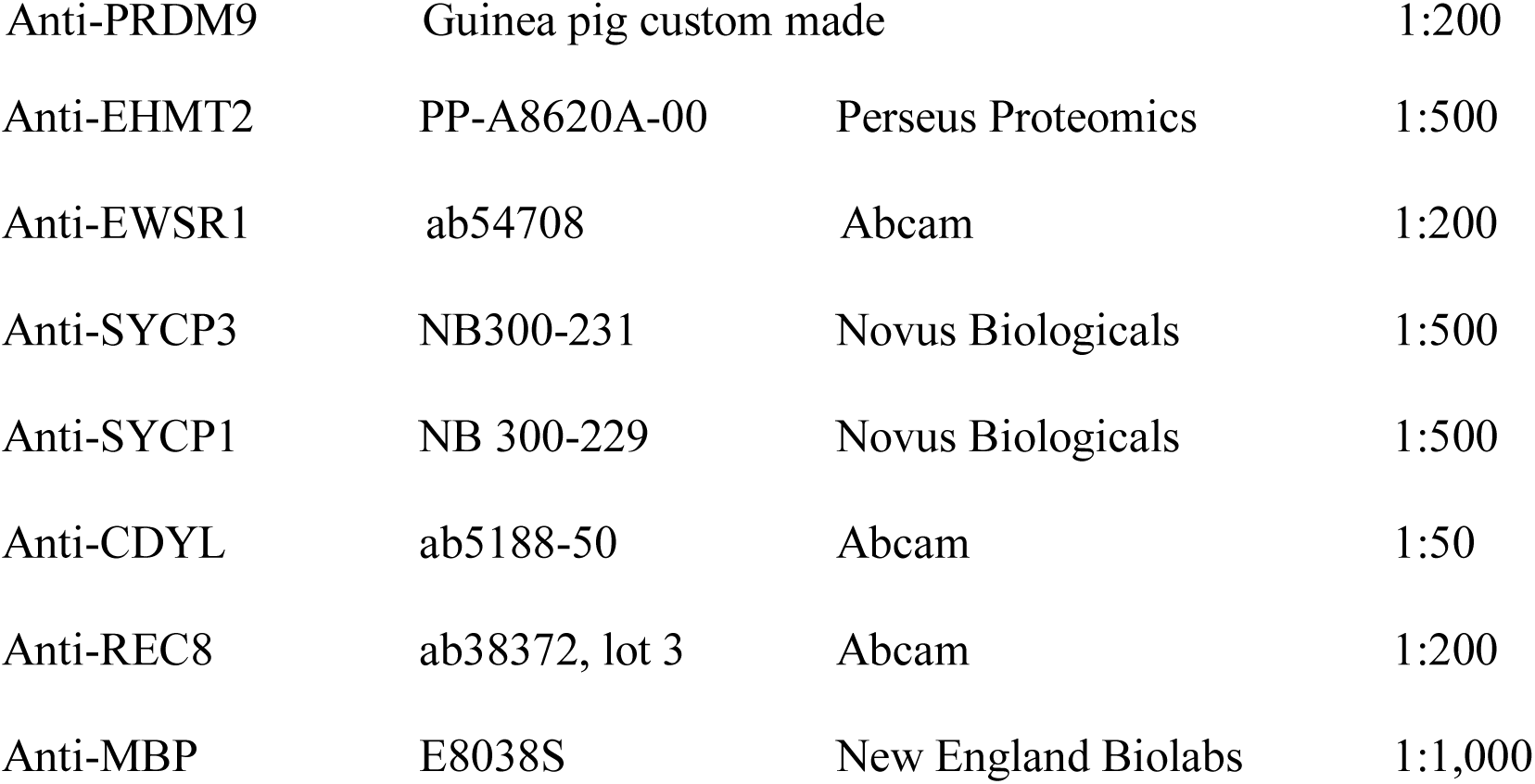

|  |  |  |
| --- | --- | --- |
| Anti-PRDM9 | Guinea pig custom made | 1:200 |

|  |  |  |  |
| --- | --- | --- | --- |
| Anti-EHMT2 | PP-A8620A-00 | Perseus Proteomics | 1:500 |
| Anti-EWSR1 | ab54708 | Abcam | 1:200 |
| Anti-SYCP3 | NB300-231 | Novus Biologicals | 1:500 |
| Anti-SYCP1 | NB 300-229 | Novus Biologicals | 1:500 |
| Anti-CDYL | ab5188-50 | Abcam | 1:50 |
| Anti-REC8 | ab38372, lot 3 | Abcam | 1:200 |
| Anti-MBP | E8038S | New England Biolabs | 1:1,000 |

Goat anti-Rabbit IgG (H+L) Secondary Antibody, Alexa Fluor^^®^^ 594 conjugate (Life Technologies, A-11037,), 1:1,000

Goat anti-Rabbit IgG (H+L) Secondary Antibody, Alexa Fluor^^®^^ 488 conjugate (Life Technologies ab150077), 1:1,000

Goat anti-Mouse IgG (H+L) Secondary Antibody, Alexa Fluor^^®^^ 488 conjugate (Life Technologies, ab150113), 1:1,000

Goat anti-Mouse IgG (H+L) Secondary Antibody, Alexa Fluor^^®^^ 647 conjugate (Life Technologies, A-21236), 1:1,000

Goat anti-Guinea pig IgG (H+L) Secondary Antibody, Alexa Fluor^^®^^ 594 conjugate (Life Technologies, A-11076), 1:1,000

Goat anti-Guinea pig IgG (H+L) Secondary Antibody, Alexa Fluor^^®^^ 488 conjugate (Life Technologies, A-11073), 1:1,000

### Yeast two-hybrid screen

The yeast two-hybrid screen was performed by co-transformation of pGBKT7-Prdm9 construct and mouse testes cDNA library cloned in pGADT7 (3×10^6^ to 10^7^ clones) (Clontech, #638848) in PJ69-4α strain. The positive interactions were selected by plating the transformants on three different selective media lacking combinations of -Trp, -Leu, -His, and -Ala, as well as supplemented with 3 mM 3-Amino-1,2,4-triazole (3-AT) for detection of more stringent interactions. Each positive clone was tested for auto activation by crossing to a yeast strain containing empty pGBKT7 plasmid. Plasmid DNA was isolated and sequenced to identify the individual cDNA library clone.

### Yeast two-hybrid screen validation and Prdm9 domain mapping

The detected open-reading frames interacting with Prdm9 were isolated and transformed in PJ69-4A strain. The shorter pGBKT7-Prdm9 constructs were transformed in PJ69-4α strain. The PJ69-4α strain was crossed with PJ69-4A strain carrying the clone of interest and plated on selective media plates for confirmation of positive interactions.

### Protein-protein *in vitro* pull-downs

Expression of PRDM9 was performed in Arctic DE3 cells. Pre-culture was grown overnight at 30°C. The cells were re-inoculated in the next day; the culture was grown for 3-4 hours and shifted for 16-24 hours at 14°C at 200 rpm. The cells were collected by centrifugation at 5,000g for 10 minutes, the pellet was ground by SPEX™ SamplePrep 6870 Freezer/Mill™ and dissolved in 1X CBB buffer (50 mM Tris-HCl, 4 mM EDTA, 200 mM sucrose, pH 7.5) +150 mM KCl, 10 ml per 1 g pellet, supplied with protease inhibitor cocktail (aprotinin, chymostatin, leupeptin, pepstatin) - Applichem, 1mM PMSF, 1mM mercaptoethanol, 0.01% NP40.

The purification of each protein was done in three steps - Sp-Sepharose, amylose beads, and FPLC on MonoS column. The pull-down of the purified PRDM9 protein and the protein of interest was done by affinity beads corresponding to the tag of the protein. The pull-down was performed by mixing 1 µg of each protein for 30’ at 34°C, followed by addition of 30 µl beads washed by K buffer (40 mM K_2_PO_4_, 10% glycerol, 0.5 mM EDTA, pH 7.5) +150 mM KCl and incubated for another 30 min. The bound proteins were eluted with SDS loading buffer.

### Co-immunoprecipitation assays

Testis material from twenty 14-dpp old C57BL/6J male mice was extracted in cold PBS, homogenized by Dounce homogenizer, passed through 40 µm cell strainer (Falcon BD # 352340), and centrifuged for 5’ at 3,000g. The pellet was re-suspended in 1 ml Pierce IP buffer (ThermoFisher Scientific, #87787) with 1 mM PMSF and 1X EDTA-free protease inhibitor cocktail (Roche). The sample was incubated for 30’ with slow rotation and centrifuged at 13,200g. For the DNAse-treated co-IP samples, 100 µl DNAse buffer and 20 U DNAse (Ambion) were added and the samples were incubated for 1 hour at room temperature. The co-IP was done by protein A or G beads (Dynabeads, Lifesciences) dependent on the antibody source. As a negative control, IgG from the same animal species was used. The extract was incubated overnight at 4°C with rotation. After washing the beads three times with 1ml Pierce IP buffer, the complexes were eluted with 200 µl GST buffer (0.2 M glycine, 0.1% SDS, 1% Tween 20, pH 2.2) for 20’ at room temperature. The sample was neutralized with 40 µl 1M Tris-HCl pH 8. For SDS-PAGE, 40 µl SDS-loading buffer was added. The samples were heated to 95°C for 5’ and subjected to electrophoresis and western blotting. Each lane contained 10 µg protein except for the input in the EHMT2 and CDYL westerns, which contained 2.5 µg to prevent overwhelming the co-IP signal.

### Chromosome spreads

For preparation of nuclear spreads from germ cells, the drying-down technique (37) was used, followed by double or consecutive immunolabeling with PRDM9/SYCP3, PRDM9/SYCP1, PRDM9/EWSR1/SYCP3, EWSR1/BRCA1/SYCP3, SYCP3/γH2AX/CREST, and PRDM9/REC8. For DNAse-treated spreads samples, slides were treated with 100 µl DNAse buffer containing 10U DNAse at 37°C for 2 h, and then followed by immunolabeling.

### PAS staining

For histological evaluation, tissues were dissected out, fixed with Bouin’s solution, embedded in paraffin wax, and 5 μm sections were prepared. Sections were stained with Periodic acid–Schiff–diastase (PAS) using standard techniques.

### Immunofluorescent staining

For protein immunolocalization, tissues were dissected out, fixed with 4% PFA solution, embedded in paraffin wax and sectioned at 5 μm. Sections were heated in a microwave in 10 mM sodium citrate buffer, pH 6.0 for 10’, and then treated with PBS containing 0.1% Triton X-100. After blocking non-specific binding sites with 10% normal donkey serum (Jackson ImmunoResearch Labs, Inc., # 017-000-121), sections were incubated with primary antibodies overnight at 4ºC and secondary antibodies for 2 h at room temperature, respectively. The slides were rinsed in PBS, stained for 3 min with 1 µg/ml DAPI (4’, 6-diamidino-2-phenylindole) (Sigma, # 28718-90-3), rinsed three times in PBS for 5’ each and mounted in Antifade reagent (Life Techonologies, # S-2828). Images were photographed with Microscope Axio Imager.Z2 (Zeiss, Germany).

### TUNEL assay

For detection of apoptosis in tissues, testis sections were subjected to fluorescence labelling of DNA strand breaks by Terminal deoxynucleotidyl transferase-mediated digoxigenin-dUTP nick-end labeling (TUNEL) assay, using the In situ cell death detection kit (Roche, # 11684795910) according to the manufacturer’s protocol. As a negative control, the TdT enzyme was omitted in parallel reactions.

## RESULTS

### Yeast two-hybrid assay identifies direct PRDM9 interactors

To search for proteins directly interacting with PRDM9, we performed a yeast two-hybrid (Y2H) screen using cloned full-length *Prdm9* as bait and a 6-month old mouse testis cDNA library as prey. Screening ~6×10^6^ colonies, we isolated a total of 329 positive clones, which after sequencing coalesced to 118 individual open reading frames. Four clones, representing C-terminal portions of *Ehmt2* (amino acids 376-1263)*, Cxxc1* (217-660), *Ewsr1* (479-655), and *Cdyl* (74-593) genes, were confirmed as interacting strongly with full-length PRDM9 by pairwise Y2H under the most discriminating conditions (Fig. 1). To identify the positions of their binding sites on the PRDM9 molecule, shorter PRDM9 fragments were cloned as bait constructs and tested by pairwise Y2H with each interacting clone (Fig. 1A and B). All four clones were found to interact with PRDM9 fragments representing the isolated KRAB domain, and the *Ehmt2* and *Ewsr1* clones interacted with the intervening region between the KRAB and PR/SET domains as well (Fig. 1B).

**Figure 1.**
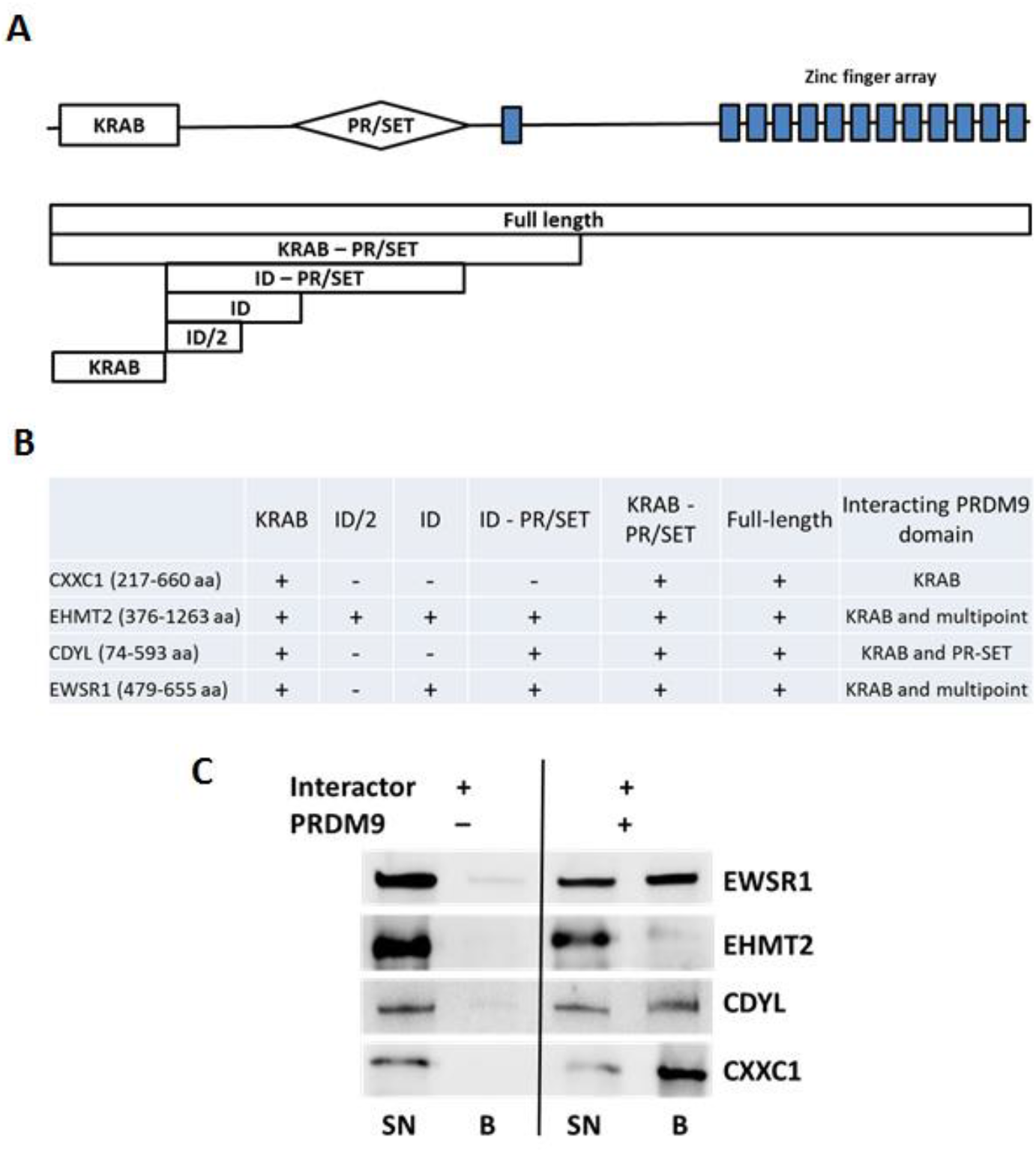
CXXC1, EWSR1, EHMT2, and CDYL directly interact with PRDM9. A. Scheme of the domain structure of PRDM9 (top) showing the positions of the KRAB, PR/SET, and the zinc finger domains. Below are schematic representations of the full-length and deletion constructs used in yeast two-hybrid assay to map the contact points of PRDM9 interacting with the other proteins. B. Results of the yeast two-hybrid assay showing that KRAB is the major protein contact domain of PRDM9. The extents of the cloned and expressed fragments of the interactor proteins are in brackets. C. Purified PRDM9 and its interactors bind to each other *in vitro.* Purified HALO-tagged EWSR1 and CDYL, or GST-tagged EHMT2 and CXXC1 were immobilized on amylose beads alone (left half), or mixed with purified full-length MBP-tagged PRDM9 (right half). Specific interactions are detected by the immobilization of interactor protein on amylose beads only in the presence of MBP-tagged PRDM9. All four proteins specifically bind to PRDM9. SN – supernatant; B – beads.

The amino acid sequences of the clones showing positive interactions with PRDM9 provide clues regarding the binding sites of these proteins. The only known functional domain in the cloned fragment of EWSR1 is a C-terminal RanBP2-type zinc finger, which has been implicated in protein binding (44). The cloned portion of CXXC1 includes acidic, basic, coiled coil domains, a Set1-interacting domain, and a PHD2 domain (48). The cloned portion of EHMT2 contains an NRSF- binding cysteine rich domain, an ankyrin domain, and a SET domain (14). The CDYL clone lacks 19 amino acids of its N-terminal chromodomain but contains the remaining part and the C-terminal ClP domain (51).

### EWSR1, EHMT2, CDYL and CXXC1 bind PRDM9 *in vitro*

Each of the interacting proteins detected by Y2H also bound to PRDM9 *in vitro*. For these tests, we cloned, expressed and purified N-terminally tagged versions of these proteins from *E. coli*. Full-length PRDM9 was cloned as a fused construct with an N-terminal MBP tag. The other four were cloned separately as fused constructs with HALO or GST tags. The tagged proteins were expressed in *E. coli* and purified. Individual proteins were mixed with PRDM9 and corresponding complexes bound to amylose beads. All four proteins bound PRDM9 *in vitro* (Fig. 1C). Interestingly, one of the two histone modifiers, EHMT2, showed relatively weaker binding to PRDM9 (Fig. 1C, second row) compared to EWSR1, CDYL and CXXC1.

### Binding of PRDM9 and its interacting proteins in spermatocytes

We next tested whether these proteins also interact with PRDM9 and with each other in mouse spermatocyte lysates by co-IP in germ cells of 14-dpp (days post-partum) juvenile mice, which are enriched for the leptotene through early pachytene stages of meiotic prophase I (Fig. 2A, left panel). The co-IP with antibodies against PRDM9 also contained EWSR1, EHMT2, and CDYL as interactors, confirming the Y2H data. *In vivo* binding of PRDM9 to EWSR1 was strong; however, its strength diminished after DNAse treatment suggesting possible bridging the interaction via DNA. In contrast, PRDM9 interactions with EHMT2 and CDYL were only slightly affected by DNAse treatment (Fig. 2A, left panel). We could not test CXXC1 for the lack of specific antibodies. We also carried out reverse co-IP experiments in which antibodies against EWSR1 pulled down PRDM9 but not EHMT2, and CDYL (Fig. 2A, central panel). Antibodies against CDYL showed co-IP with PRDM9 and EHMT2, which was retained after DNAse treatment, but failed to show any signal with EWSR1 (Fig. 2A, right panel). The available antibodies against EHMT2 were not suitable for co-IP.

**Figure 2.**
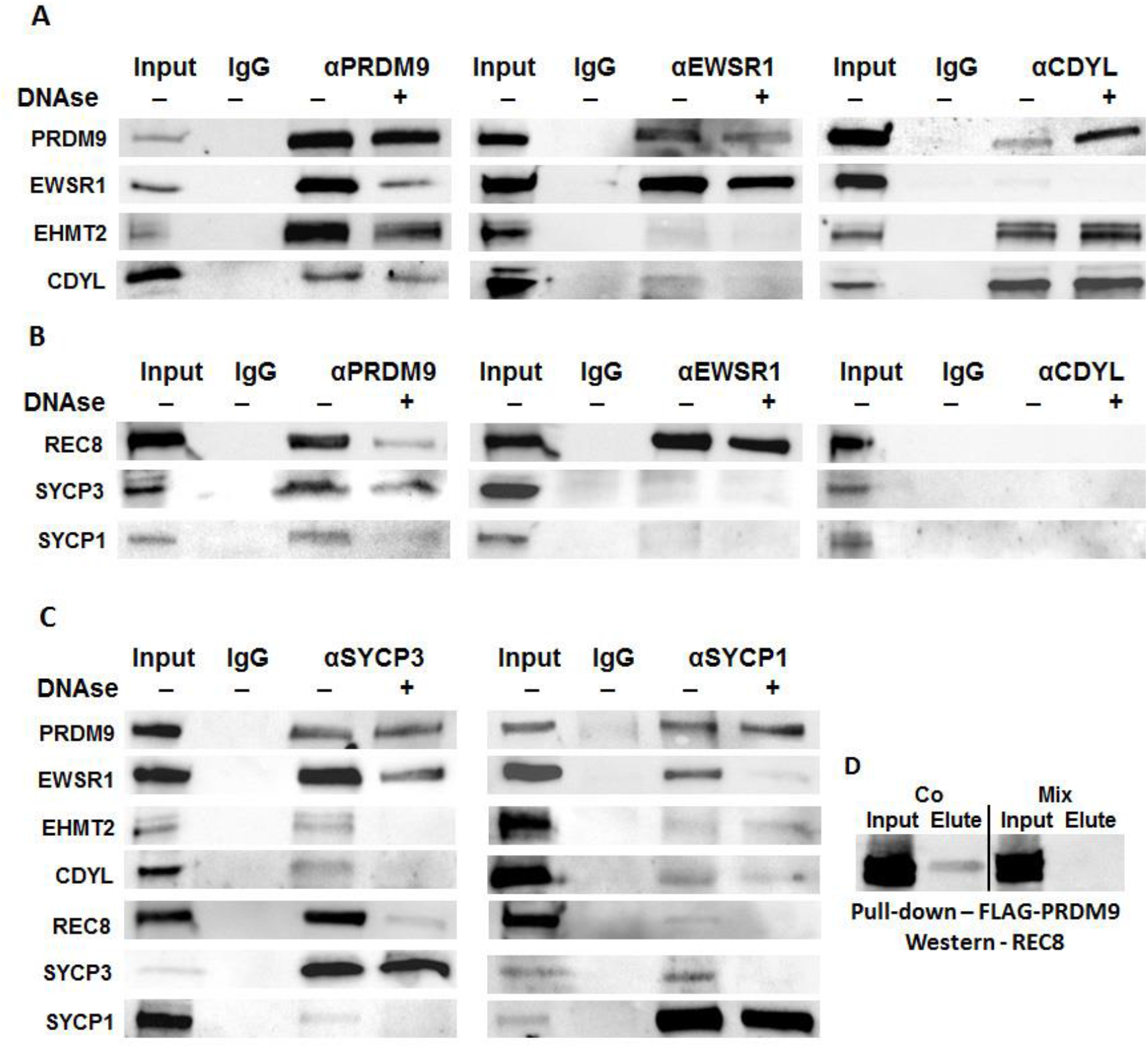
PRDM9, EWSR1, and CDYL co-immunoprecipitate each other and chromosomal axis/synaptonemal complex proteins other from wild type 14-dpp spermatocytes. A. PRDM9 and its interactors co-IP each other. Left panel, Co-IP with anti-PRDM9; central panel, co-IP with anti-EWSR1; right panel, co-IP with CDYL. In each panel, lane 1, Input – 10 µg (2.5 µg in the CDYL and EHMT2 blots); lane 2, IgG – 10 µg co-IP with non-immune IgG; αPRDM9, αEWSR1, αCDYL – 10 µg co-IP with the respective antibody, either non-treated (lane 3), or DNAse-treated (lane 4). The antibodies used to detect specific proteins on western blots are shown on the left. B. PRDM9 and its interactors also interact with chromosomal axis/SC proteins. Co-IP with the same antibodies used in A, but probed with antibodies against chromosome axis/SC proteins as marked on the left. C. Reciprocal co-IP shows that SC proteins co-IP PRDM9 and its interactors. Co-IP with SYCP3 (left panel) or SYCP1 (right panel). The antibodies used for detection on western blots are on the left. D. Interaction between REC8 and PRDM9 in cultured cells. REC8 was cloned under a V5 tag, and PRDM9 was cloned under a FLAG-tag. The two proteins were either co-expressed or separately expressed in HEK 293 cells. Specific interaction between the two proteins was detected in extracts of cells where the two proteins were co-expressed (Co, left) but not in mixed extracts of cells where the two proteins were expressed separately (Mix, right). Note that the interacting band is the middle of three bands indicating that it is a phosphorylated form.

We conclude that PRDM9 forms a separate complex with EWSR1 *in vivo*, independent of its binding with EHMT2 and CDYL. We also found evidence of strong interaction between EHMT2 and CDYL *in vivo.* Taken together, these data suggest the possible existence of both a PRDM9-EWSR1 complex, and a separate PRDM9-EHMT2-CDYL complex.

### Interactions with the synaptonemal complex

Because PRDM9-bound hotspots are translocated to the chromosomal axis where DSBs are subsequently formed and repaired, we asked whether PRDM9 and its interacting proteins bind components of the chromosomal axis, and found this to be the case. Specifically, we tested for co-IP with the meiotic-specific cohesin REC8, chromosomal axis/SC lateral element protein SYCP3, and the SC central element protein SYCP1. Co-IP with anti-PRDM9 pulled down REC8, SYCP3, and SYCP1, and these interactions were partially retained after DNAse treatment (Fig. 2B, left panel).

Antibodies against EWSR1 showed a strong co-IP signal with REC8, which was retained after DNAse treatment, but only a faint signal with SYCP3 and SYCP1 (Fig. 2B, central panel). The interaction with REC8 appears to be with the phosphorylated form of the protein as only antibodies able to detect the phosphorylated form of REC8 with apparent electrophoretic mobility corresponding to ~85 kDa (17) showed positive signal with PRDM9 and EWSR1. Co-IP with antibodies against CDYL did not detect interaction with any of these proteins (Fig. 2B, right panel).

In reverse, antibodies against the lateral element protein SYCP3 strongly pulled down PRDM9 and EWSR1 but showed weak signal with EHMT2 and CDYL, which was not retained after DNAse treatment (Fig. 2C, left panel). They also pulled down REC8 and SYCP1, but in a DNAse sensitive manner, confirming that DNA is indeed essential for the integrity of the chromosomal axis. Antibodies against the central element protein SYCP1 pulled down PRDM9, an interaction retained after DNAse treatment. They also strongly pulled down EWSR1 but the signal was sensitive to DNAse treatment (Fig. 2C, right panel). Weak signals were also detected with EHMT2 and CDYL. Unfortunately, all available antibodies against REC8 were not suitable for co-IP.

Although these data provide strong evidence that PRDM9 and EWSR1 associate with the meiotic-specific cohesins *in vivo*, we were unable to detect direct interaction between PRDM9 and REC8, by Y2H or *in vitro* after expression of the proteins in *E. coli* (data not shown). Consequently, to provide additional confirmation of interactions with the chromosomal axis/SC *in vivo*, we used a HEK293 mammalian cell expression system and showed co-IP of the phosphorylated form of REC8 with PRDM9 when the two were co-expressed in the same cells (the middle band in Fig. 2D, second lane), but not when they were expressed separately and the cells mixed before extraction. The lack of interaction between the purified proteins *in vitro* could therefore be explained by the lack of REC8 phosphorylation when expressed in *E.coli*. In addition, this interaction may require another protein intermediate such as EWSR1 which is naturally present in HEK293 cells.

### PRDM9 binding to other proteins is dependent on its own binding to DNA

To investigate whether PRDM9 binding to other proteins is dependent on its zinc finger domain and the ability to bind to DNA, we created a new PRDM9 functional knockout mouse model, B6(Cg)-Prdm9^tm3.1Kpgn^/Kpgn (designated here as *Prdm9*^*tm3.1Kpgn*^). In this mouse, we placed a mutation creating an alternative splice acceptor site 44 bp inside the 5’-end of exon 12 that codes for the entire zinc finger domain, causing a frame shift with a stop codon. As a result, the mutant protein contains amino acids 1-381 of PRDM9 but lacks all of its zinc fingers (Fig. 3A) and the ability to bind DNA at specific sequences (49). It is stable and appears as a 48-kDa band on Western blots in testes from both heterozygous and homozygous mutant animals (Fig. 3B). Both male and female mice homozygous for this mutation are sterile. Spermatogonia and spermatocytes were found in mutant testes but no post-meiotic spermatids were observed (Fig. 3C). Cytological staining of spreads of mutant spermatocytes show cells arrested at an aberrant pachytene-like stage, with inappropriate γH2AX staining on autosomes, and asynapsis of homologous chromosomes (Fig. 3D). Lack of the PRDM9 zinc finger domain also led to increased apoptosis during meiosis compared to wild type (Fig. 3E, *p* < 0.05). This phenotype is very similar to the one described in *Prdm9*^*TM1Ymat*^ KO mice which have no PRDM9 protein (21, 45).

**Figure 3.**
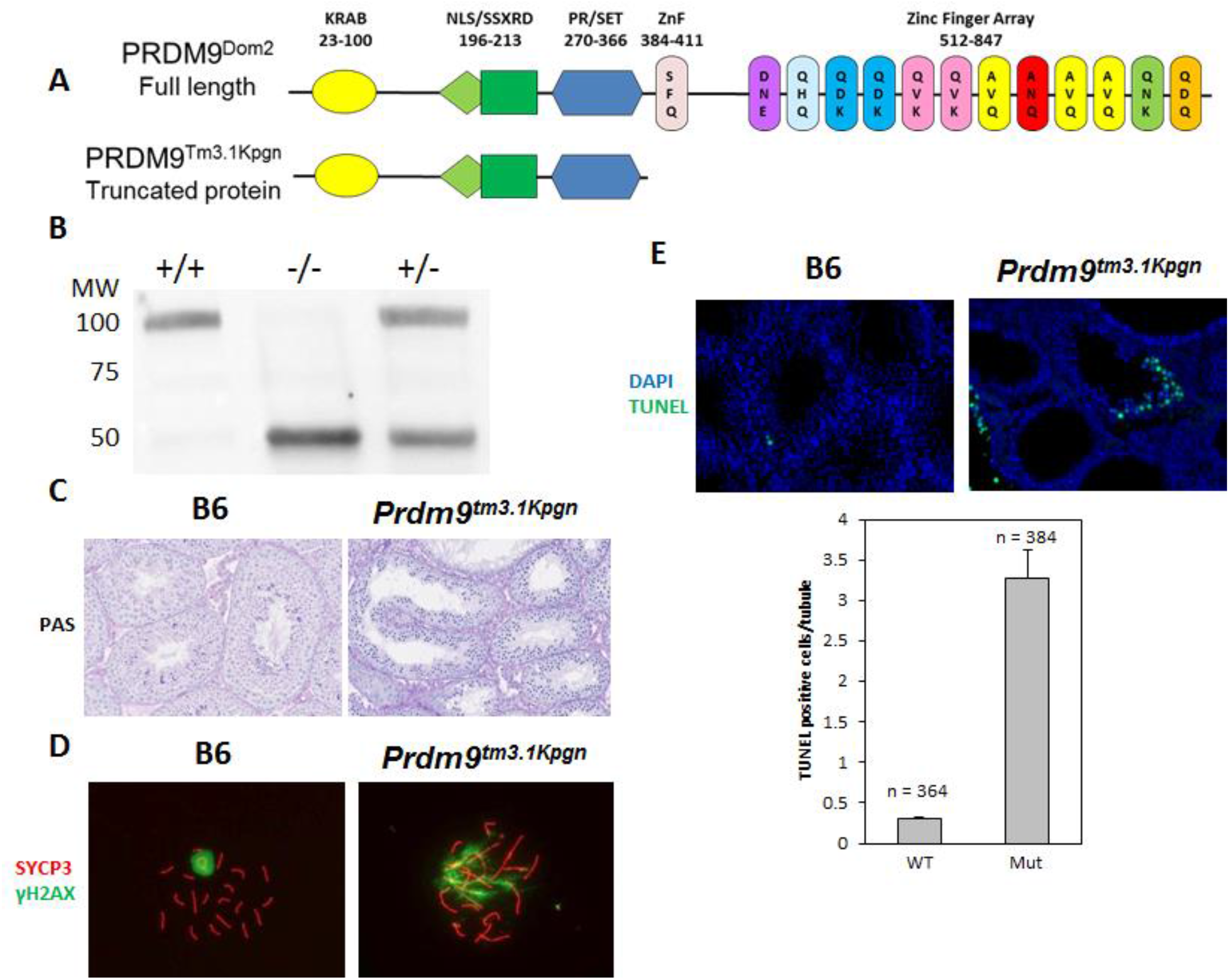
Characterization of the *Prdm9*^*tm3.1Kpgn*^ mutant. A. Schematic representation of wild type PRDM9 and mutant PRDM9^Tm3.1Kpgn^ protein. B. Western blotting showing that the truncated protein is expressed in both homozygous and heterozygous mice. The bands correspond to the predicted molecular mass of 98 kDa for the wild type and 48 kDa for the truncated protein. The amount of the two forms in the heterozygous testes is approximately equal. Testis extract used: +/+, wild type B6, -/-, homozygous mutant, +/-, heterozygous. C. PAS staining of testis tubule sections from wild type (left) and mutant mice (right). Note the lack of postmeiotic cells in the mutant indicating that the germ cells undergo meiotic arrest. D. SYCP3/γH2AX/CREST staining on germ cell spreads showing a pachynema cell in the wild type (left), with a γH2AX signal (green) restricted only to the sex body, and the latest stage found in the mutant, representing late zygonema – pachynema transition. E. Increased levels of apoptosis in the mutant compared to the wild type. Upper panel, TUNEL staining in wild type (left) and mutant testis (right). Bottom panel, quantitation of the apoptotic cells in wild type and mutant testis. Asterisk, *p* < 0.05.

The truncated 48 kDa version of PRDM9 was immunoprecipitated from testis lysates using anti-PRDM9 antibodies directed against the remaining portion of the molecule (Fig. 4A, left panel). These co-IP experiments showed weak and inconsistent signals with most of the proteins that associate with full length PRDM9 in three independent experiments. Consistent weak interactions were detected only with EWSR1 and SYCP3 (Fig. 4A, left panel). These results suggest that, although EWSR1, CDYL, and EHMT2 can directly interact with the isolated N-terminal region as well as the full length protein *in vitro*, their *in vivo* interactions require PRDM9 binding to DNA.

**Figure 4.**
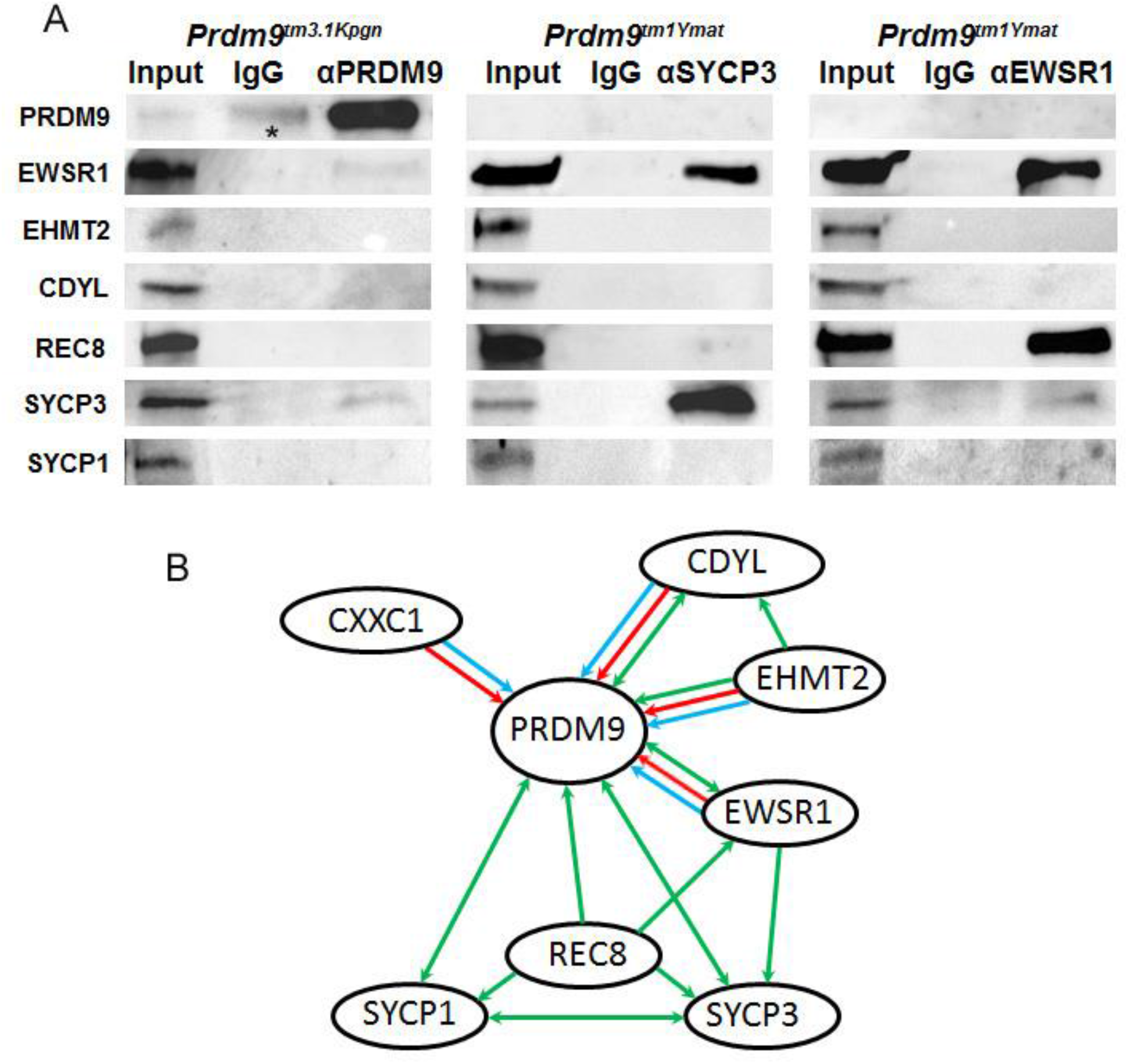
Protein-protein interactions in 14-dpp spermatocytes. A. Interactions in spermatocytes of PRDM9 mutant mice. Left panel: co-IP with anti-PRDM9 in testes of *Prdm9*^*tm3.1Kpgn*^ mice lacking its DNA-binding ZnF domain. The asterisk indicates overlapping signal from the heavy chain of IgG. All protein amounts are as in Fig. 2. Central panel: co-IP with anti-SYCP3 in testes of *Prdm9*^*tm1Ymat*^ mice lacking PRDM9 protein. Right panel: co-IP with anti-EWSR1 in testes of *Prdm9*^*tm1Ymat*^ mice. In all panels, lane 1 – input, lane 2 – IgG, lane 3 – co-IP. B. Summary of protein-protein interactions detected by all methods. Blue line – direct interactions found by yeast two-hybrid assay; red line – direct interactions detected by mixing purified proteins and isolating their complexes *in vitro*; green line – interactions detected by co-IP. The arrows show the direction of interactions detected. Double-headed arrows show interactions confirmed by reciprocal co-IP.

### EWSR1 binds to the chromosomal axis in the absence of PRDM9

To determine if binding of EWSR1, EHMT2 and CDYL to the chromosomal axis depends on their association with PRDM9 or whether these proteins can bind independently, we performed co-IP with anti-SYCP3 antibodies in testes of *Prdm9*^*tm1Ymat*^ knockout mice (21) that do not express any PRDM9 protein (45). EWSR1 showed a strong positive signal (Fig. 4A, central panel), indicating that it can bind to the synaptonemal complex independently of PRDM9. In contrast, CDYL and EHMT2 were not pulled down by SYCP3 in the absence of PRDM9. In addition, the lack of REC8 and SYCP1 signal indicates chromosomal axis impairment. Together with the co-IP data from wild type testis, this indicates that the CDYL-EHMT2 complex associates with the chromosomal axis only in the presence of PRDM9 bound to hotspot DNA.

Co-IP with EWSR1 in *Prdm9*^*tm1Ymat*^ knockout mice showed a strong signal with REC8 and a weak signal with SYCP3. EHMT2 and CDYL were not detected as EWSR1 interactors (Fig. 4A, right panel). These results suggest that EWSR1 can provide a link between PRDM9 and the chromosomal axis through the meiotic-specific cohesin complexes containing REC8.

A summary of all protein-protein interactions detected is presented in Fig. 4B.

### PRDM9 colocalizes with its interactors in mid- to late zygonema

Our previous work established that PRDM9 is only present during meiosis of in pre-leptonema, leptonema, and zygonema, a period of roughly 48 hours (45). To confirm that PRDM9 is co-expressed in the same cells and co-localizes within nuclei with its interactors, we performed double staining with combinations of antibodies against PRDM9 and its interactors in seminiferous tubules and spermatocyte spreads. We first determined the spatial and temporal co-localization of PRDM9 and EWSR1 in seminiferous tubules of 14-dpp juvenile mice (Fig. 5A, top panel). EWSR1 showed high expression in spermatogonia (Fig. 5A, Fig. 5C, top panel, arrowhead) and Sertoli cells (marked by GATA4, Fig. 5C, arrow) located at the base of seminiferous tubule, and remained present in the nuclei of spermatocytes located in the tubule lumen (Fig 5A, top panel, arrow). At 14-dpp, when most spermatocytes are in leptonema and zygonema (45), PRDM9 and EWSR1 are clearly co-expressed in those cells (Fig. 5A, top panel, arrow). Because the EWSR1 signal is weaker in PRDM9-positive cells, we sought to confirm that these proteins co-localize in spermatocyte spreads of 14-dpp mice by performing triple staining with EWSR1, PRDM9, and SYCP3, using the chromosomal axis protein SYCP3 as a marker of meiotic progression (Fig. 5B). PRDM9 and EWSR1 were clearly co-expressed in pre-leptonema to mid-zygonema nuclei. The EWSR1 signal increased dramatically in pachynema, a time when PRDM9 has disappeared from the germ cell nuclei. Interestingly, the EWSR1 signal was excluded from the sex body in pachynema as demonstrated by its lack of co-localization with BRCA1 (Fig. 5B, yellow arrow).

**Figure 5.**
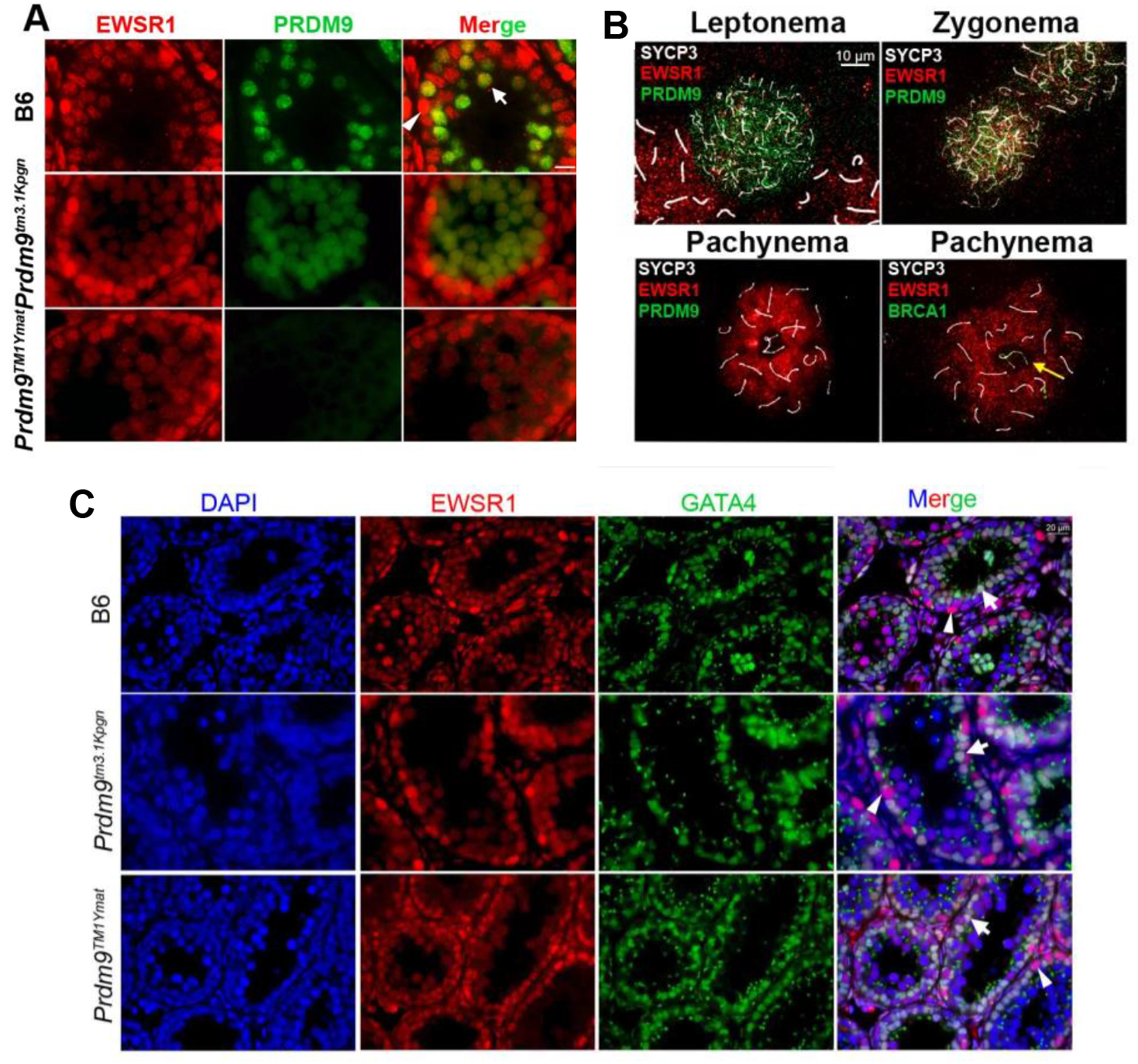
EWSR1 is co-expressed with PRDM9 in early meiotic prophase. A. Immunofluorescence analysis of EWSR1 (red) – PRDM9 (green) co-expression in tissue sections from testis tubules of 14-dpp mice. Top panel: wild type B6 mice. Middle panel: *Prdm9*^*tm3.1Kpgn*^. Arrowhead, spermatogonia with high EWSR1 expression. Arrow, spermatocytes with EWSR1 and PRDM9 positive signals. Lower panel: *Prdm9*^*tm1Ymat*^. B. Co-localization analysis of EWSR1 and PRDM9 in spermatocyte spreads. Triple staining for EWSR1 (red), SYCP3 (white), and PRDM9 (green) in leptonema, zygonema (top panels) and pachynema (lower left panel) or EWSR1 (red), SYCP3 (white), and BRCA1 (green) in pachynema (lower right panel) of B6 mice. EWSR1 is excluded from the sex body marked by BRCA1 in pachynema (yellow arrow). C. Double staining with EWSR1 and the Sertoli cell marker GATA4. EWSR1 expression is stronger in spermatogonia (arrowhead) than in Sertoli cells (arrow). Top panels: wild type B6 mice; middle panels: *Prdm9*^*tm3.1Kpgn*^; lower panels: *Prdm9*^*tm1Ymat*^. EWSR1 shows similar expression pattern in mutants and in wild type.

In *Prdm9*^*tm3.1Kpgn*^ mice lacking the PRDM9 zinc finger domain, only a diffuse EWSR1-PRDM9 co-localization pattern is seen in seminiferous tubules (Fig. 5A, second row). This mutant undergoes meiotic arrest around the zygonema-pachynema transition and the seminiferous tubule lumens contains very few of the EWSR1-positive, PRDM9-negative pachynema germ cells. Very similar EWSR1 staining is found in *Prdm9*^*tm1Ymat*^ KO mice which do not express any PRDM9 protein (Fig. 5A, third row).

We conclude that EWSR1 and PRDM9 are co-expressed in early meiotic prophase for the entire time of PRDM9 expression.

EHMT2 and CDYL signals were not detectable on spermatocyte spreads. For this reason, we tested for co-expression of PRDM9, CDYL, and EHMT2 in seminiferous tubules of 14-dpp mice.

In spermatocytes, CDYL staining in the nucleus persists only until leptonema, after which it is translocated to the cytoplasm. Strong nuclear and cytoplasmic CDYL signal was detected in some cells close to the basal membrane which were PRDM9 negative (Fig. 6, top row, arrowhead). Weak nuclear CDYL signals were detected in germ cells that also showed weak PRDM9 signal (Fig. 6, top row, open arrow). We consider these cells to be spermatocytes in pre-leptonema or leptonema based on their tubule staging and PRDM9 positivity (45). The increase of PRDM9 signal was accompanied by translocation of the CDYL signal to the cytoplasm (Fig. 6, top row and short arrows in second row,). These cells are most probably in leptonema to zygonema. Pachynema cells abundant in the lumen showed strong cytoplasmic CDYL but not PRDM9 signals (Fig. 6, third row, long arrow).

**Figure 6.**
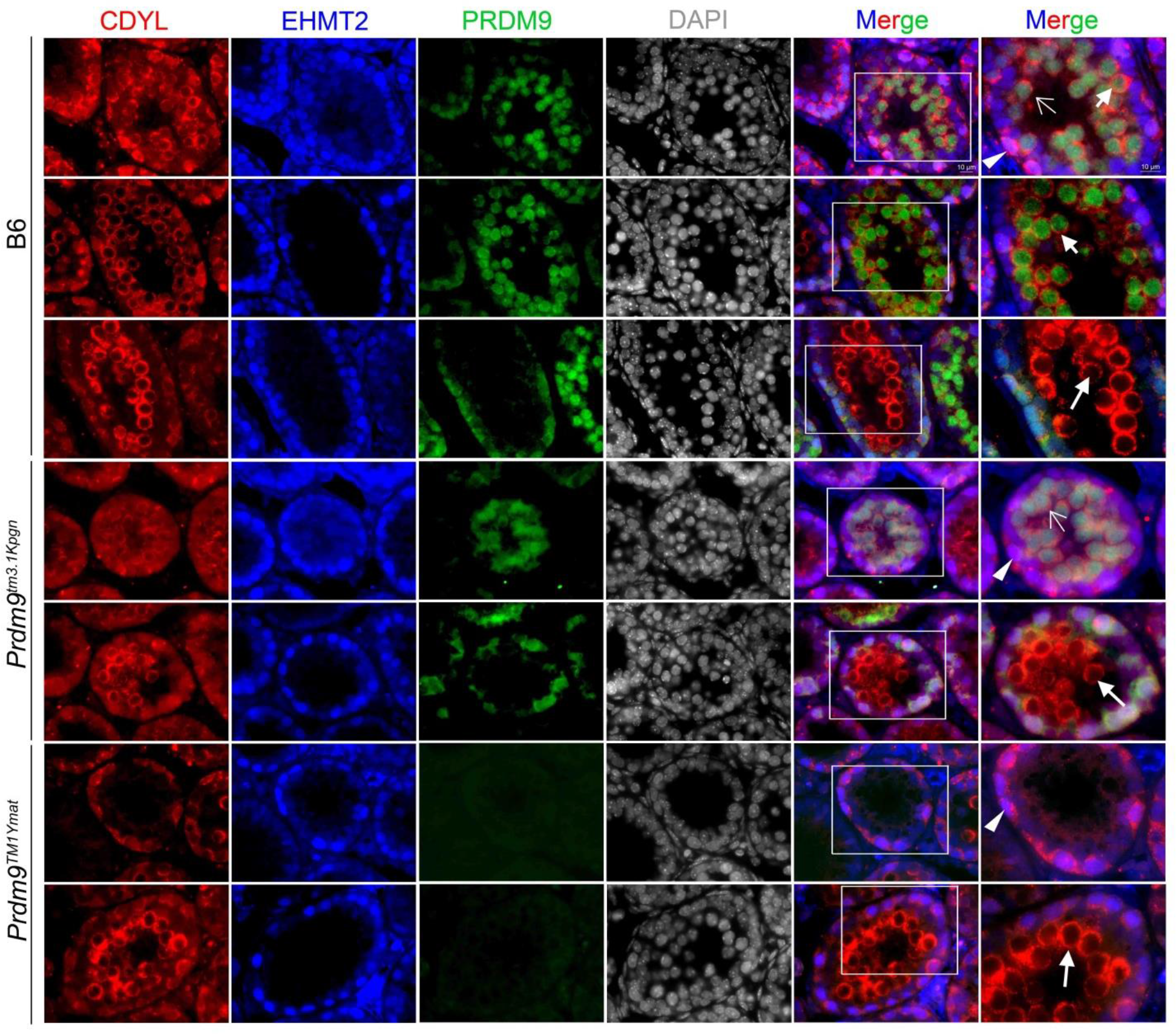
CDYL and EHMT2 are co-expressed with PRDM9 in early meiotic prophase. Co-expression of CDYL (red), EHMT2 (blue), and PRDM9 (green) expression in tissue sections from testis tubules of 14-dpp mice. Top three rows, co-expression analysis in B6 mice. Fourth and fifth rows, expression in *Prdm9*^*tm3.1Kpgn*^ mutant. Sixth and seventh row, expression in *Prdm9*^*tm1Ymat*^ mutant. Arrowhead, Sertoli cells showing strong nuclear and cytoplasmic CDYL and strong EHMT2 signal PRDM9 negative. Open arrow, pre-leptonema to leptonema cells with weak nuclear CDYL, EHMT2, and PRDM9 signals. Short arrow, late leptonema to early zygonema cells with strong PRDM9, cytoplasmic CDYL, and lack of EHMT2 signals. Long arrow, pachynema cells with strong cytoplasmic CDYL but lack of EHMT2 and PRDM9 signals.

Neither the presence of truncated PRDM9 nor the complete loss of PRDM9 affected the localization patterns of CDYL from pre-leptonema through zygonema/early pachynema-like stages (Fig 6). Triple staining in testes of *Prdm9*^*tm3.1Kpgn*^ and *Prdm9*^*tm1Ymat*^ mutant mice confirmed the meiotic arrest phenotype found by EWSR1-PRDM9 staining. *Prdm9*^*tm3.1Kpgn*^ testes showed CDYL -PRDM9-positive cells extending to the lumen, and appearance of a few CDYL cytoplasmic-positive, PRDM9-EHMT2-negative cells (Fig. 6, fourth and fifth rows). Similarly, *Prdm9*^*tm1Ymat*^ testes which lack PRDM9 protein contained some CDYL cytoplasmic-positive cells (Fig. 6, sixth and seventh rows).

EHMT2 had a nuclear pattern of expression very similar to CDYL but, unlike CDYL which was translocated to the cytoplasm, disappeared entirely from the cells in late leptonema – early zygonema. Strong EHMT2-positive, but PRDM9 and GATA4 negative spermatogonia (Fig. 6, top row, arrowhead, and Fig. 7, arrowhead) were found close to the basal membrane. Sertoli cells, marked by the presence of GATA4, showed weaker EHMT2 signal (Fig. 7, arrow). Weak EHMT2 and weak PRDM9 signals were detected in germ cells attached to the basal membrane (Fig. 6, top row, open arrow) representing spermatocytes in pre-leptonema or leptonema. Cells in zygonema were characterized by an increase of the PRDM9 signal and disappearance of the EHMT2 signal (Fig. 6, top row and short arrows in second row,). Pachynema cells abundant in the lumen had neither EHMT2 nor PRDM9 signals (Fig. 6, third row, long arrow). The localization of EHMT2 in *Prdm9*^*tm3.1Kpgn*^ and *Prdm9*^*tm1Ymat*^ mutant testes followed the same pattern (Fig. 6, fourth to seventh rows).

**Figure 7.**
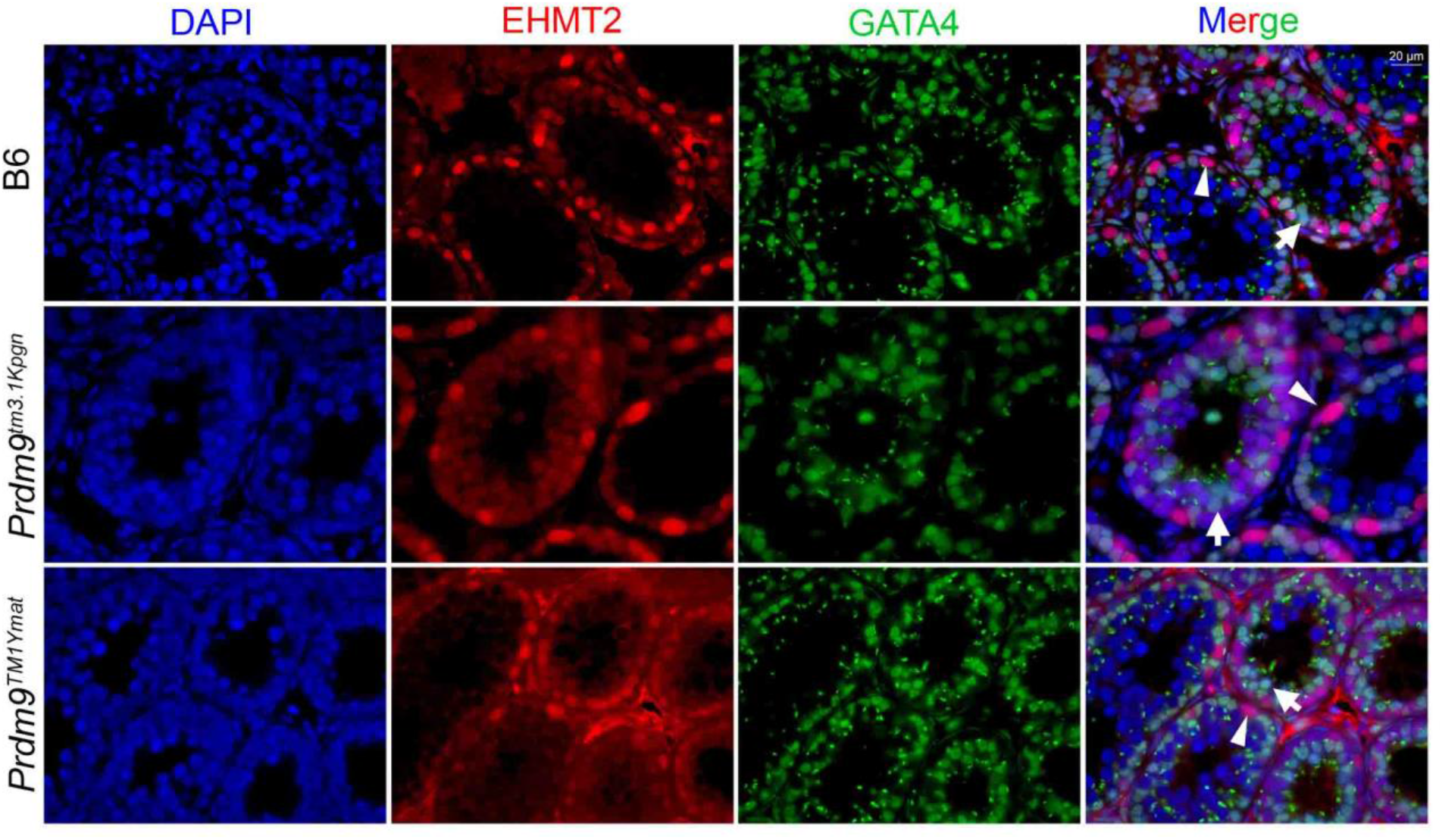
EHMT2 and GATA1 are co-expressed in Sertoli cells. EHMT2 expression is stronger in spermatogonia (arrowhead) compared to Sertoli cells (arrow). Top panel: wild type B6 mice; middle panel: *Prdm9*^*tm3.1Kpgn*^; lower panel: *Prdm9*^*tm1Ymat*^. EHMT2 shows similar expression pattern in mutants and in wild type.

It appears that EWSR1, CDYL, and EHMT2 coincide with each other and with PRDM9 only in pre-leptonema to early zygonema, when all four of them are present at low concentrations. After that, CDYL is translocated to the cytoplasm; EHMT2 disappears altogether, and both EWSR1 and PRDM9 increase in expression until the zygonema-pachynema transition. At pachynema, PRDM9 disappears whereas EWSR1 expression dramatically increases. These findings suggest a temporal order in the formation of the PRDM9-bound complexes. The PRDM9-CDYL-EHMT2 complex is restricted to pre-leptonema and leptonema, after which it is dissolved with the disappearance of CDYL and EHMT2 from the nucleus. The PRDM9-EWSR1 complex is probably formed later in leptonema and persists until late zygonema until PRDM9 disappears from the nucleus.

### Meiotic progression into zygonema requires PRDM9-bound hotspot translocation to the chromosomal axis

Because the co-IP evidence suggested that PRDM9 binds to meiosis-specific cohesins such as REC8, we sought to determine whether co-localization between and PRDM9 and REC8 could be detected in early meiotic prophase on spermatocyte spreads which can be staged by the appearance of the REC8 signal (24). In pre-leptonema, REC8 staining shows both diffuse and punctate patterns. This is similar to the pattern of PRDM9 at the same stage. In leptonema, REC8 starts forming rod-like structures (Fig. 8A, top row) which become predominant in zygonema as elements of the chromosomal axes (Fig. 8A, second row). The co-localization of these axis structures with some of the PRDM9 foci at late zygonema was apparent (Fig. 8A, third row, arrows). *Prdm9*^*tm3.1Kpgn*^ testes had a REC8-PRDM9 pattern similar to wild type zygonema but showed less evidence for co-localization (Fig. 8A, fourth row, arrows). The PRDM9 signal in this mutant has a more diffuse and less punctate pattern than in wild type spermatocytes (Fig. 8A, fourth row; see also Fig. 8B and 8C, fourth rows) suggesting that the punctate pattern reflects PRDM9 binding to hotspot DNA. *Prdm9*^*tm1Ymat*^ testes lacking PRDM9 contained mostly cells with REC8 patterns reminiscent of pre-leptonema to early zygonema (Fig. 8A, fifth row), indicating that progression of spermatocytes into zygonema requires an intact PRDM9-bound hotspot translocation to the chromosomal axis.

**Figure 8.**
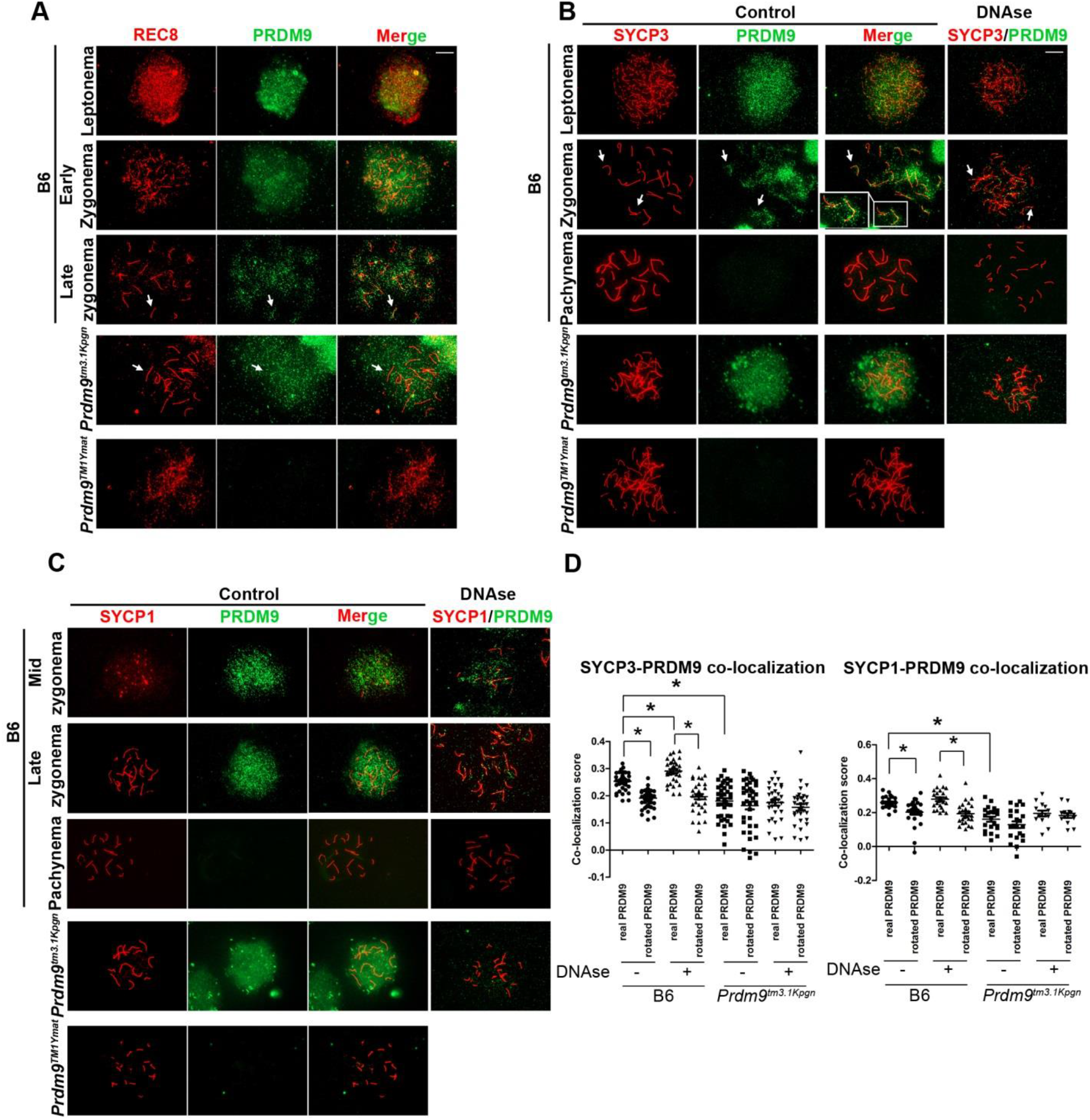
REC8, SYCP3, and SYCP1 co-localize with PRDM9 in early meiotic prophase. A. Co-localization of REC8 (red) and PRDM9 (green) in spermatocyte spreads. First three rows, B6 spermatocytes in leptonema (top row), early zygonema (second row), and late zygonema (third row). Arrows show co-localization pattern of the two proteins becoming more prominent at later stages. Fourth row, co-localization pattern in late zygonema-like cells in the *Prdm9*^*tm3.1Kpgn*^ mutant. Note the more diffuse pattern of the truncated PRDM9 lacking its DNA-binding domain compared to wild type in the third row. Fifth row, leptonema or early zygonema-like cell in the *Prdm9*^*tm1Ymat*^ mutant. B. Co-localization of SYCP3 (red) and PRDM9 (green) in spermatocyte spreads. Disposition of the panels is the same as in A. Arrows show co-localization pattern of the two proteins. C. Co-localization of SYCP1 (red) and PRDM9 (green) in spermatocyte spreads. Disposition of the panels is the same as in A. D. Quantitation of SYCP3/PRDM9 (left panel) and SYCP1/PRDM9 (right panel) co-localization in B6 and *Prdm9*^*tm3.1Kpgn*^ mutant compared to the co-localization pattern of images where the two signals were inverted to 180° relative to each other. Asterisk, *p* <0.05.

### PRDM9 binding to DNA promotes translocation to the chromosomal axis

The co-IP experiments also suggested that PRDM9 binds to SC proteins. For this reason, we tested whether PRDM9 co-localizes with SC proteins in early meiotic prophase by double staining with a combination of PRDM9 and either SYCP3 or SYCP1 antibody on spermatocyte spreads. SYCP3 and SYCP1 show punctate or rod-like staining in early meiosis, similar to that of REC8 (Fig. 8B and 8C). Both SC protein signals overlapped with PRDM9, with a clearer pattern in late zygonema, when a significant part of the diffuse PRDM9 signal disappears (Fig. 8B and 8C, second rows, arrows). Because this pattern makes it difficult to determine whether the proteins truly co-localize, we performed a DNAse treatment which removed the loop DNA and resulted in loss of most of the PRM9 signal not associated with the chromosomal axis (Fig. 8B and 8C, fourth columns). Under these conditions, the association of PRDM9 with both SC proteins was apparent (Fig. 8B and 8C, second rows, fourth columns, arrows). To determine whether this association was statistically significant, we compared the co-localization of actual signals to images in which one of the signals is inverted 180^O^ relative to the other (25). In wild type mice, PRDM9 showed co-localization with SYCP3 in zygotene (Li’s ICQ value p = 6×10^−5^, Fig. 8D, left panel). PRDM9 also showed co-localization with SYCP1 when the latter first appeared in mid-zygotene (p = 2.2×10^−5^, Fig. 5D, right panel). The statistical significance of the difference between actual and inverted images was even greater after DNAse treatment (Fig. 8D, Li’s ICQ value p = 1.33×10^−12^ for SYCP3 and p = 6×10^−14^ for SYCP1.).

Importantly, we compared these results with the colocalization pattern of the *Prdm9*^*tm3.1Kpgn*^ mutant. At best, we found very weak evidence of significant co-localization between PRDM9 and either SYCP3 or SYCP1 in this mutant (Fig. 8D, left panel, p=0.13, and Fig. 8D, right panel, p=0.16). The lack of co-localization was confirmed after DNAse treatment (Fig.8B and 8C, fourth rows, fourth columns, and Fig. 8D). The difference in the colocalization patters between wild type and mutant spermatocytes was statistically significant for both combinations (p<0.05).

The combined co-IP and cytological evidence indicates that the binding of PRDM9 to DNA promotes the translocation of DNA-PRDM9-protein complexes to the chromosomal axis.

## DISCUSSION

### PRDM9 directly interacts with EWSR1, CXXC1, EHMT2, and CDYL

Yeast two-hybrid screens and *in vitro* binding assays show that EWSR1, CXXC1, EHMT2, and CDYL can directly bind to PRDM9 via its KRAB domain as well as by additional contact points extending further to the PR/SET domain (Fig. 1). With the exception of CXXC1, for which specific antibodies are not currently available, we confirmed that these interactions also occur in mouse spermatocytes (Fig. 2 and 4). However, the spatial and temporal pattern of expression of each of these proteins *in vivo* suggests that these interactions probably occur at different stages of meiotic progression and involve two distinct PRDM9 complexes, one with EHMT2 and CDYL and another with EWSR1 and possibly CXXC1. Several lines of evidence show that PRDM9 forms dimers at hotspots (3, 36), suggesting that PRDM9 can use its KRAB domain to bind two proteins at the same time, with PRDM9-EHMT2-CDYL, as one complex, and PRDM9-EWSR1-CXXC1, as another.

EHMT2 and CDYL are expressed strongly in both Sertoli cells and spermatogonia, before PRDM9 appears at the onset of meiosis, and they remain present in the subsequent pre-leptotene and leptotene stages when PRDM9 is first detected in the nuclei of germ cells (45). In early zygonema, EHMT2 disappears from the nucleus, and CDYL is translocated out of the nucleus and remains in the cytoplasm into pachynema (Fig. 6). CDYL interacts strongly with EHMT2 by co-IP (Fig. 2), suggesting that their interaction takes place in any cell type where they are present together – Sertoli cells, spermatogonia, and early spermatocytes. However, the two proteins can each bind to PRDM9 *in vitro*. Thus, their similar co-IP patterns with PRDM9 in spermatocytes could reflect both their binding to each other and independently to PRDM9. CDYL also complexes EHMT2 in mouse embryonic stem cells, but there the two proteins are found bound together in heterochromatin regions, including the inactivated X chromosome in females (15). This is in marked contrast to spermatocytes where PRDM9 binding to DNA results in chromatin activation by catalyzing H3K4 and H3K36 trimethylation of nearby nucleosomes (4, 38), suggesting that a triple PRDM9-CDYL-EHMT2 complex may well have a different molecular function than the double CDYL-EHMT2 complex lacking PRDM9.

EWSR1 expression coincides with PRDM9 in leptonema and zygonema. These data, together with the results of Y2H, *in vitro* binding, and the strong mutual co-IP of the two proteins, make a strong case that the two proteins physically interact with each other from leptonema to late zygonema. However, unlike PRDM9, EHMT2 and CDYL, EWSR1 dramatically increases in nuclei at pachynema (Fig. 5B). The continued, increased expression of EWSR1 at later stages suggests that EWSR1 likely provides two distinct functions: an earlier one when complexed with PRDM9 and a later one when PRDM9 has disappeared. In humans EWSR1 interacts with BARD1 (43), which apart from other recombination factors (BRCA1, RAD51 etc.) interacts with SETDB1 (19) – a H3K9 histone methyltransferase, which in turn interacts with CDYL-EHMT2 (31). PRDM9, which also has a SET domain, may play role equivalent to that of SETDB1 by taking central place as a shared binding partner of EWSR1 and CDYL-EHMT2. This possibility is further supported by the recent findings that PRDM9, like SETDB1, is capable of trimethylating H3K9 *in vitro* (38, 50), although the presence of H3K9ac at hotspots (11) likely prevents this activity *in vivo*.

CXXC1 binds directly to PRDM9 in Y2H and *in vitro*. Unfortunately, the lack of appropriate antibodies prevented us from detecting its interactions *in vivo*. What is known is that in mammals CXXC1 is part of the Set1 complex responsible for most of H3K4 trimethylation in somatic cells (27, 40). In ES cells it is required for both H3K4me3 deposition after DNA damage and the subsequent acetylation of H3K9 at the same nucleosomes (13). With respect to its possible functions in meiosis, studies of Spp1, the yeast homolog of CXXC1, provide clues to its possible role in mammalian meiosis. In *S. cerevisiae*, DSBs initiate at H3K4me3 sites found near promoters (9). DSB formation at these sites is promoted when they become tethered to the chromosomal axis by Spp1/CXXC1 (1, 42). We now show evidence that CXXC1 binds to PRDM9 directly, and that REC8 interaction with PRDM9 is indirect, possibly requiring mediation by EWSR1. This suggests that CXXC1 may act cooperatively with EWSR1, providing an additional link between PRDM9-bound H3K4-trimethylated sites and the chromosomal axis to promote proper DSB formation and repair.

### PRDM9 interacts and colocalizes with meiotic-specific cohesin REC8 and synaptonemal complex proteins SYCP1 and SYCP3

We found no evidence for direct binding between PRDM9 and the meiotic cohesin REC8 in either our Y2H screen or when the two proteins were tested *in vitro* after expression in *E. coli.* However, the two proteins showed ample evidence of interaction in spermatocytes both by pulldown experiments (Fig. 2A) and cytological co-localization (Fig. 8A), and when they were expressed together in HEK 293 cell cultures. In both cases, the shifted mobility of REC8 and the specificity of the antibody we used, suggest that only phosphorylated REC8 interacts with PRDM9. In addition, the substantial reduction in co-IP after DNAse treatment suggests that these proteins require an additional molecule to provide a link which needs not to be meiosis-specific. Our co-IP results suggest EWSR1 as a likely candidate; it is a generally “sticky” protein (39) that strongly pulls down both PRDM9 and REC8 in spermatocytes independently of the presence of DNA; the other likely candidate is CXXC1, given its known role in bring hotspots to the chromosome axis in yeast, and these are not mutually exclusive possibilities.

These findings opens the possibility that complexes including PRDM9, EWSR1, and other proteins, such as CXXC1, play a role in homolog recognition by bringing the homologous hotspot DNA sequences bound to PRDM9 in contact with cohesins and subsequently to the chromosomal axis.

Our findings that the synaptonemal complex proteins REC8 and SYCP3, and to some extent SYCP1, not only interact with each other, but also with PRDM9, and EWSR1, support the concept that these proteins play important roles in bringing activated hotspots from out in the DNA loops down to the chromosome axis, stabilizing hotspot location there, and then participating in DSB formation and repair. Interestingly, both SYCP3 and SYCP1 showed stronger binding with PRDM9 than with each other (Fig. 2C), given that SYCP1 and SYCP3 do not bind directly to each other but through SYCP2. SYCP1 participates in the formation of the SC central element interacting with intermediates such as SYCP2, SYCE1, SYCE2 and SYCE3, and TEX12 (see (8) for review). This raises the possibility that PRDM9 contact with SYCP1 has a function beyond bringing hotspot DNA to the chromosomal axis.

Judging by the properties of its yeast homolog Spp1, CXXC1 may well function in combination with the histone modifications placed by PRDM9 at hotspots in the transport of hotspots to the chromosome axis. Once DSBs have been formed, EWSR1 is known to promote ssDNA invasion into dsDNA, forming meiotic Holliday Junctions (20, 39), and has been shown to contribute to DSB repair (29). And EHMT2 and CDYL are known to be involved in the establishment of closed chromatin states (15, 28, 52) such as those evidenced by the presence of γH2AX around as yet unrepaired meiotic DNA lesions (12). However, we never detected any presence of closed chromatin marks such as H3K9me2/me3 near active hotspots. Instead, nucleosomes in the immediate vicinity of DSB sites are modified to create an open chromatin state required for repair (4, 52). In this regard, the temporary presence of EHMT2-CDYL at PRDM9-bound complexes could help restrict the extent of H3K4me3-marked open chromatin to ensure proper space for the subsequent repair before DSBs are initiated by SPO11.

### Protein interactions are dependent of PRDM9 binding to DNA

Our characterization of protein-protein interactions in the *Prdm9*^*tm3.1Kpgn*^ mutant, which lacks the zinc finger domain, showed that PRDM9 binding to various partners is dependent on the presence of this domain. This seems to be in contrast with the Y2H data showing that PRDM9 binding to EWSR1, EHMT2, CDYL, and CXXC1 occurs through its N-terminal part including the KRAB and PR/SET domains. A likely explanation of this apparent contradiction is that EWSR1, CDYL, and EHMT2 may be predominantly bound to DNA and/or chromatin in the nuclei, whereas PRDM9 can be present in both the nuclear matrix and bound to hotspot DNA, in which case, the interactions can only occur in the context of DNA packed in chromatin. In addition, each PRDM9 fragment in the Y2H assay is bound to DNA through the GAL4 DNA-binding domain of the vector which may provide the conditions for proper binding.

### EWSR1, but not EHMT2 and CDYL, bind to the SC in the absence of PRDM9

PRDM9 presence is not necessary for binding of EWSR1 to the chromosomal axis, but is required for binding of EHMT2-CDYL complexes to the chromosomal axis as evidenced by our SYCP3 co-IP results in testes of *Prdm9*^*tm1Ymat*^ KO mutant (Fig. 4, central panel). EWSR1 binding to SYCP3 and REC8 in this mutant is as strong as in wild type (Fig. 2B and 4) suggesting that it is associated with the chromosomal axis from the earlier stages of its formation.

### Model of events occurring before recombination initiation

Our results show extensive protein-DNA and protein-protein interactions as part of recombination-related events in early meiotic prophase. Our data suggest the following working model for the mechanisms and dynamics of these events (Fig. 9). In early leptonema, a fraction of the PRDM9 molecules present in the nucleus bind to hotspots as dimers in which only one subunit binds to DNA (3). Both subunits participate in the trimethylation of histone 3 at lysine-4 and lysine-36 ensuring the deposition of H3K4me3 and H3K36me3 on both sides of the PRDM9 binding site. The KRAB domains of the PRDM9 subunits bind CDYL and EHMT2, restricting the extent of trimethylation to 2-4 nucleosomes on each side (4, 38) (Fig. 9A). By the end of leptonema, CDYL and EHMT2 are removed from the complex and the nucleus, and the hotspot-bound PRDM9 and the adjacent nucleosomes bind EWSR1 and CXXC1 (Fig. 9B). These complexes then bind to REC8 and translocate the hotspot DNA from out in chromosome loops down to the chromosomal axis where REC8 integrates into the axis, with the participation of SYCP3 (Fig. 9C). PRDM9 remains bound to the hotspots until DSB are initiated by SPO11, meanwhile coming into contact with SYCP1 (Fig. 9D). PRDM9 then disappears from the nucleus in late zygonema, first the unbound molecules, and later the ones associated with the SC, whereas EWSR1 further participates in the formation and resolution of Holliday junctions at pachynema, and carries out additional functions.

**Figure 9.**
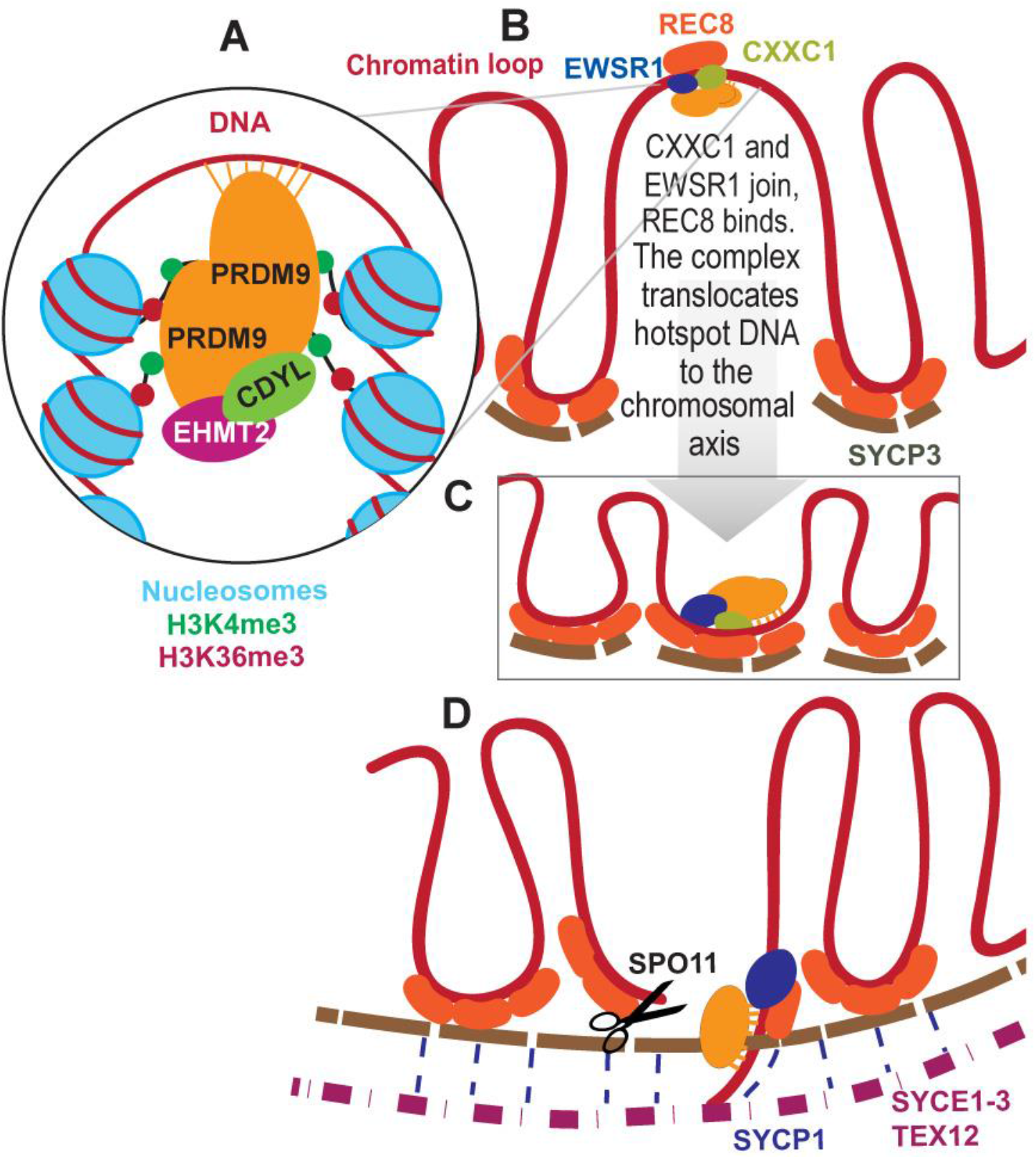
Model of events occurring before recombination initiation (see text for details).

As interesting as this model may be in suggesting new experimental directions, it does lack at least one essential element: what brings the homologous hotspot down to the chromosome axis for precise pairing and DNA exchange. We know from previous studies (2) that precise recombination can occur between homologous hotspots even when one has lost the ability to bind PRDM9, removing PRDM9 binding at the homolog or its consequences as the requisite signal. The model proposed here brings one homolog down to the chromosome axis, but we still lack a model for what brings its homologous partner into apposition.

## ACKNOWLEDGMENTS

The authors thank Anita Hawkins for technical help, Mary Ann Handel for critical reading and helpful suggestions, Neil Hunter for advice on immunofluorescence, and Attila Toth for information about HORMAD proteins.

